# PLB-985 neutrophil-like cells as a model to study *Aspergillus fumigatus* pathogenesis

**DOI:** 10.1101/2021.07.28.454178

**Authors:** Muhammad Rafiq, Flora Rivieccio, Ann-Kathrin Zimmermann, Corissa Visser, Alexander Bruch, Thomas Krüger, Katherine González Rojas, Olaf Kniemeyer, Matthew G. Blango, Axel A. Brakhage

## Abstract

Fungal infections remain a major global concern. Emerging fungal pathogens and increasing rates of resistance mean that additional research efforts and resources must be allocated to advancing our understanding of fungal pathogenesis and developing new therapeutic interventions. Neutrophilic granulocytes are a major cell type involved in protection against the important fungal pathogen *Aspergillus fumigatus*, where they employ numerous defense mechanisms, including production of antimicrobial extracellular vesicles. A major draw-back to work with neutrophils is the lack of a suitable cell line system for the study of fungal pathogenesis. To address this problem, we assessed the feasibility of using differentiated PLB-985 neutrophil-like cells as an *in vitro* model to study *A. fumigatus* infection. We find that dimethylformamide-differentiated PLB-985 cells provide a useful recapitulation of many aspects of *A. fumigatus* interactions with primary human polymorphonuclear leukocytes. We show that differentiated PLB-985 cells phagocytose fungal conidia and acidify conidia-containing phagolysosomes similar to primary neutrophils, release neutrophil extracellular traps, and also produce antifungal extracellular vesicles in response to infection. In addition, we provide an improved method for the isolation of extracellular vesicles produced during infection by employing a size-exclusion chromatography-based approach. Advanced LC-MS/MS proteomics revealed an enrichment of extracellular vesicle marker proteins and a decrease of cytoplasmic proteins in extracellular vesicles isolated using this improved method. Ultimately, we find that differentiated PLB-985 cells can serve as a genetically tractable model to study many aspects of *A. fumigatus* pathogenesis.

**IMPORTANCE:** Polymorphonuclear leukocytes are an important defense against human fungal pathogens, yet our model systems to study this group of cells remains very limited in scope. In this study, we established that differentiated PLB-985 cells can serve as a model to recapitulate several important aspects of human polymorphonuclear leukocyte interactions with the important human fungal pathogen *Aspergillus fumigatus*. The proposed addition of a cultured neutrophil-like cell line to the experimental toolbox to study fungal pathogenesis will allow for a more mechanistic description of neutrophil antifungal biology. In addition, the easier handling of the cell line compared to primary human neutrophils allowed us to use PLB-985 cells to provide an improved method for isolation of neutrophil-derived extracellular vesicles using size-exclusion chromatography. Together, these results provide significant tools and a baseline knowledge for the future study of neutrophil-derived extracellular vesicles in the laboratory.

## INTRODUCTION

Fungal infections remain a tremendous source of global morbidity and mortality. More than 1 billion individuals are affected by fungal infections per year, with invasive infections killing numbers comparable to other leading bacterial pathogens (>1.5 million per year; gaffi.org; (1)). Deadly invasive infections are caused by a relatively small number of fungi, with most of these attributed to members of the genera *Candida, Pneumocystis, Cryptococcus*, and *Aspergillus* (2). *Aspergillus fumigatus* is the major cause of aspergillosis and is particularly dangerous to immunocompromised individuals suffering from neutropenia (3). There is also emerging evidence to suggest that invasive aspergillosis may contribute to COVID-19-related deaths (4, 5), but challenges in safely obtaining bronchoalveolar lavage samples from these patients have often made confirmatory diagnosis difficult. Despite the obvious importance of *A. fumigatus* in the clinic, our understanding of this important pathogen remains lacking in many aspects, in part due to a lack of tractable experimental systems in the laboratory.

Mammals are continuously challenged by fungal pathogens. In fact, asexual spores of *A. fumigatus*, termed conidia, are thought to be inhaled by humans on a scale of hundreds per day (6). For most fungi, the body temperature of mammals is too high to allow for growth, but for organisms like *A. fumigatus* that thrive in compost piles at high temperatures, the human host provides fertile ground in the absence of a functional immune system (7). However, humans do have multiple additional defenses, including a mucociliary escalator to remove particles from the lungs, a robust epithelium, resident alveolar macrophages that eliminate the majority of the remaining fungal conidia, and infiltrating polymorphonuclear leukocytes (PMNs) that aid in clearance of conidia and destruction of fungal hyphae, among others (3, 8). Neutrophils play an essential role in antifungal defense, due to their importance in killing fungal hyphae. This is well-illustrated by the high susceptibility of neutropenic patients to *A. fumigatus* infections in the clinics (reviewed in (3, 9)).

Studies in primary human neutrophils have revealed the capacity of these cells to phagocytose conidia and release granules, neutrophil extracellular traps (NETs), and extracellular vesicles in response to invading pathogens (10-12). Phagocytosis occurs in conjunction with recognition of pathogen-associated molecular patterns by host pathogen recognition receptors. Studies in zebrafish and mice have shown that these internalized conidia can even be passed from neutrophils to macrophages for destruction of the fungus (13), implying complex intracellular trafficking. In neutrophils and macrophages however, internalized wild-type *A. fumigatus* conidia are capable of stalling phagolysosomal acidification to facilitate outgrowth.

In addition to phagocytosis, NETs are an important mechanism of defense against *A. fumigatus* that are produced in response to fungal recognition in a CD11b-dependent manner (11, 14). NET production is most abundant against hyphae, but NETs are also sometimes produced in response to resting and swollen conidia (15). In vivo, NETs were shown to be present in mouse lungs during infection but were generally dispensable for fungal clearance (15, 16), suggesting that alternative measures are required to eliminate fungal hyphae that escaped phagocytosis as conidia.

Neutrophilic granules and extracellular vesicles are two heterogeneous subcellular populations that also contribute widely to the antimicrobial response. Proteins like lactoferrin associated with granules are known to inhibit *A. fumigatus* conidial germination due to iron sequestration (17) and the abundant granule protein myeloperoxidase inhibits fungal hyphal growth (18). Extracellular vesicles were also shown to provide an antifungal mechanism against *A. fumigatus* hyphae and conidia (10). Antifungal extracellular vesicles produced by neutrophils in response to infection are capable of associating with the fungal cell wall and are in some cases internalized to deliver an unknown antifungal cargo. Intriguingly, the antifungal effect of these extracellular vesicles appeared to be tailored to the infecting pathogen, as spontaneously released vesicles and vesicles released in response to infection with a knockout strain deficient in production of the conidial pigment dihydroxynaphthalene (DHN)-melanin were not antifungal against wild-type fungus (10).

The dissection of the molecular mechanisms behind these differences in antifungal potential are difficult to discern in the absence of a tractable model system for the genetic manipulation of neutrophils. Despite the importance of neutrophils in antifungal defense, the majority of studies are still performed in primary human PMNs isolated from venous blood, a resource that can only be used for small-scale studies and is unfortunately highly variable between blood donors. A laborious isolation procedure and limited half-life of about 19 hours *ex vivo* (19) also make more mechanistic studies difficult. For all these reasons, establishment of a cell culture line to study *A. fumigatus* pathogenesis in neutrophils or neutrophil-like cells would be highly advantageous. In recent years some options have emerged to investigate specific aspects of neutrophil biology in culture (20). In particular, the myeloid cell line PLB-985, a derivative of HL-60 granulocytic cells, has recently emerged as a tractable model to study NET formation in response to bacterial infection after differentiation into neutrophil-like cells (21, 22). Of note, the related HL-60 cell line has also been used previously to deliver azole drugs to *A. fumigatus*-infected mice (23), but not to our knowledge in studies of *A. fumigatus* pathogenesis.

In this study we hypothesized that differentiated human PLB-985 neutrophil-like cells could serve as model system to study the host-pathogenesis of *A. fumigatus* in a blood donor-independent manner. We tested this hypothesis by defining the phagocytosis and intracellular trafficking of *A. fumigatus* conidia by PLB-985 cells, the production of neutrophil extracellular traps, and the production of extracellular vesicles produced in response to fungal infection. We find numerous similarities between this cell line and primary neutrophils that suggest differentiated PLB-985 cells can serve as a tractable *in vitro* model system for further study of some aspects of *A. fumigatus* pathogenesis.

## RESULTS

### Differentiated PLB-985 neutrophil-like cells phagocytose opsonized *A. fumigatus* conidia

We set out to assess the feasibility of using *N,N-*dimethylformamide (DMF)-differentiated PLB-985 (dPLB) cells as a model for neutrophil phagocytosis, NET production, and extracellular vesicles release in response to *A. fumigatus* infection. Upon infection, neutrophils are known to rapidly phagocytose and process *A. fumigatus* conidia. We therefore challenged dPLB cells with opsonized *A. fumigatus* resting conidia and assessed various parameters of infection.

First, we determined the phagocytic capacity of PLB-985 cells using confocal laser scanning microscopy following incubation of cells with fluorescein isothiocyanate (FITC)-labelled, opsonized conidia (green). We observed that dPLBs were capable of phagocytosis, as evidenced by a phagosomal membrane surrounding the conidia (**Fig. 1A**). A similar result was confirmed for primary human PMNs with this experimental setup (**Fig. S1A**). Interestingly, dPLB cells only infrequently phagocytosed unopsonized conidia, whereas primary human neutrophils indiscriminately phagocytosed both unopsonized and opsonized conidia (data not shown). To better quantify phagocytosis, we analyzed 5,000 host cells post-infection by single-cell analysis using imaging flow cytometry. For this we counterstained with calcofluor white (CFW; blue) to elucidate non-phagocytosed conidia (**Fig. 1B and Fig. S1B to C**). Quantification of imaging flow cytometry data revealed that approximately 22% of wild-type and 26% of non-pigmented Δ*pksP* conidia were phagocytosed by dPLBs after 2 hours, while undifferentiated PLB-985 cells exhibited limited phagocytosis of both wild-type and non-pigmented conidia (**Fig. 1C**). As expected, these phagocytosis percentages were lower than those of primary human PMNs, which phagocytosed 62% of wild-type and 54% of Δ*pksP* conidia after 2 hours and 90% of wild-type and 85% of Δ*pksP* conidia after 4 hours of infection (**Fig. S1D**). We also measured the induction of cell damage following infection using lactate dehydrogenase release assays and found levels below the limit of detection for dPLB cells after 2 and 4 hours of incubation (data not shown). To determine if infection also resulted in an activated immune response, we measured proinflammatory cytokine levels by enzyme-linked immunosorbent assay (ELISA). Co-incubation of dPLB cells with wild-type spores resulted in production of significantly increased levels of interleukin (IL)-8, but not IL-1β after 24 hours of co-incubation, comparable to primary neutrophils at the corresponding earlier time points (**Fig. 1D and Fig. S2**). Together, these results suggested that DMF-differentiated PLB-985 cells could serve as a suitable model for *A. fumigatus* pathogenesis and warranted further investigation.

**FIG 1.**
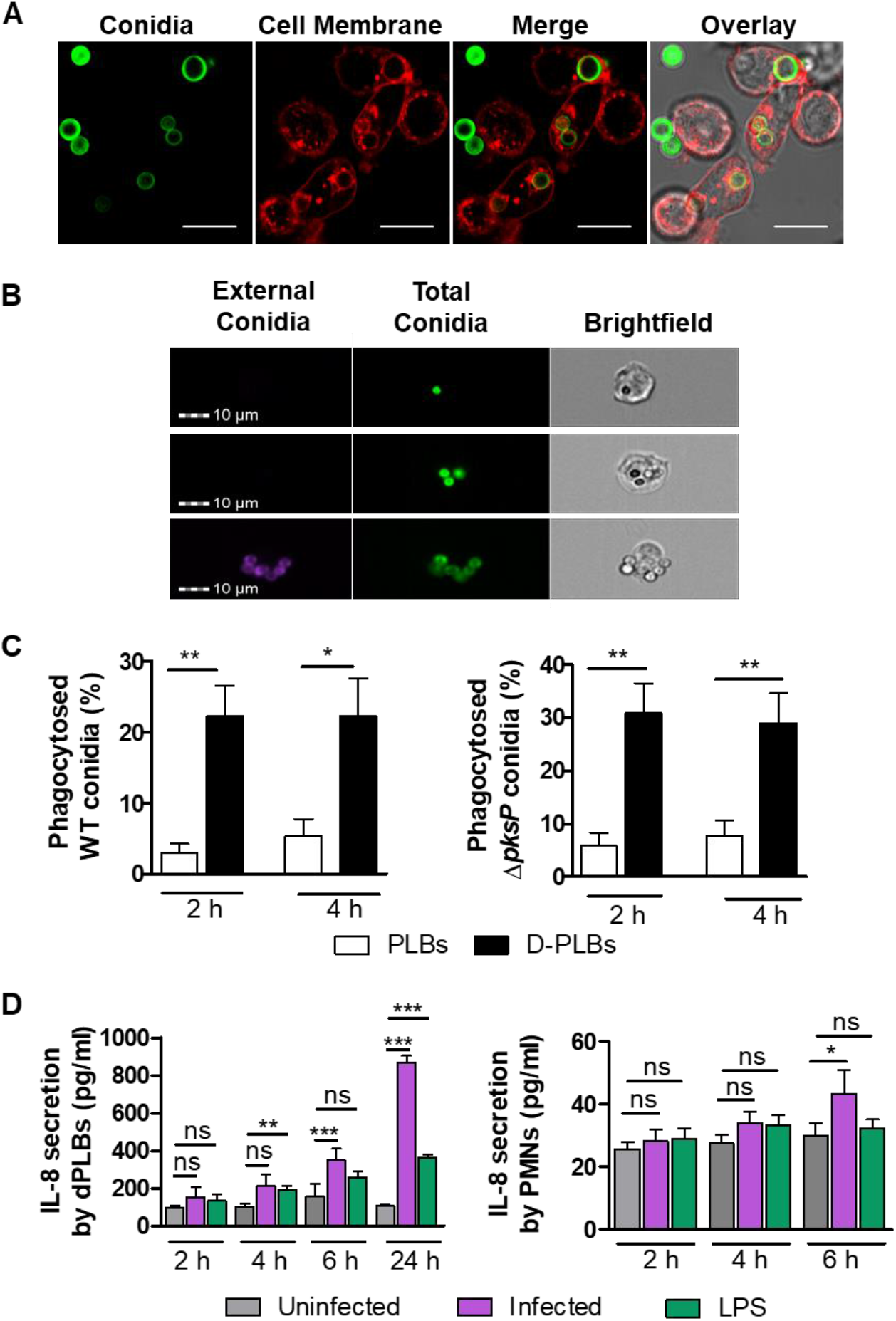
Phagocytosis of *A. fumigatus* conidia by PLB-985 cells. **(A)**. Confocal microscopy images of wild-type conidia (FITC-stained; green) phagocytosed by dPLB cells after 4 hours co-incubation. Membranes were stained with Cell Mask (red). Images are representative of three independent biological experiments. Scale bars are 10 µm. **(B)** Representative images of wild-type FITC-stained conidia phagocytosed by dPLB cells after 4 hours co-infection obtained by imaging flow cytometry (top two rows). External conidia are visualized by counter staining with CFW (purple), as seen in the bottom row. Images are representative of three independent experiments. **(C)** Quantification of PLB-985 cell phagocytosis using the IDEAs software on 5,000 cells per condition with either wild-type or Δ*pksP* conidia from four independent experiments **(D)** ELISA detection of IL-8 cytokine released from infected dPLBs and primary human PMNs at different time points. Lipopolysaccharide (LPS) was included for comparison of a bacterial stimulus. Data are presented as mean ±SD, from six biological replicates; * = p < 0.05.

### Internalized *A. fumigatus* conidia are processed inside dPLB phagolysosomes

Following internalization of conidia by professional phagocytes like macrophages and neutrophils, conidia-containing phagosomes fuse with lysosomes to acidify the compartment and aid in fungal killing. Numerous additional enzymes are activated upon phagolysosomal acidification to degrade and digest the internalized fungal conidia. The acidification and maturation of phagolysosomes are regulated by protein complexes assembled on the phagolysosomal membrane, including lysosomal-associated membrane protein (LAMP)-1, 2, and 3; several vacuolar ATPases (V-ATPases); Ras-related protein (RAB) 5 and 7; Flotillin 1 and 2; and numerous others (24). DHN-melanin on the surface of conidia can interfere with these processes in alveolar macrophages, monocytes, and primary neutrophils to delay fungal processing and phagolysosomal acidification (25). To test whether dPLB cells process conidia similarly to primary human PMNs, we first stained the cells with LAMP-1, a general endocytic marker on the membrane of phagolysosomes. LAMP-1 showed a clear signal around phagolysosomes of infected cells (**Fig. 2A**), and the intensity of the signal did not differ between wild-type and *ΔpksP* conidia (**Fig. 2B**). Furthermore, loading of dPLB cells with Lysotracker, a weak base that becomes fluorescent under acidic conditions, showed that the *ΔpksP* conidia*-*containing phagosomes were significantly more acidified than those containing wild-type conidia, revealing that DHN-melanin can also block the acidification process in the dPLB model as well (**Fig. 2C to D**).

**FIG 2.**
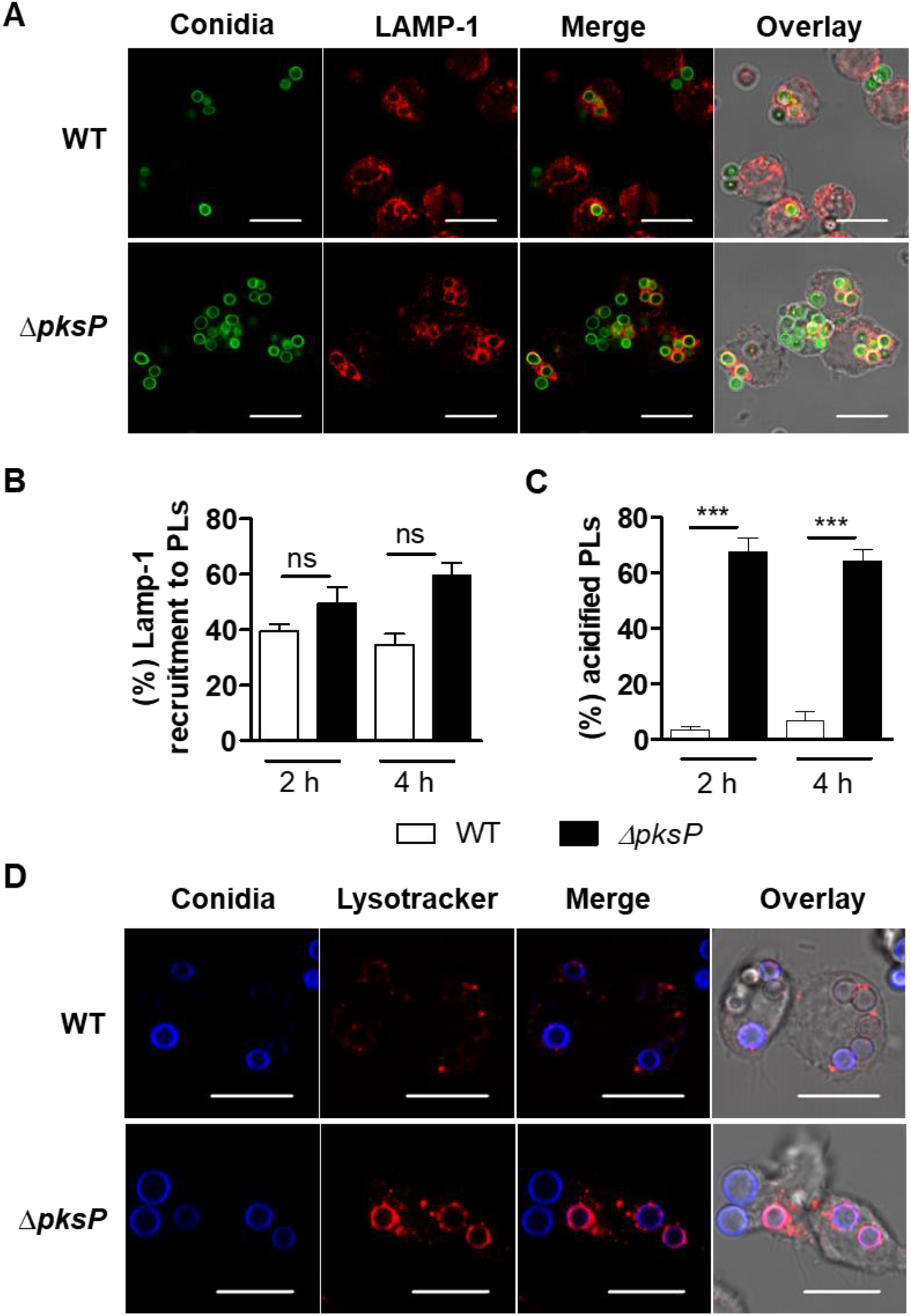
Processing of *A. fumigatus* conidia inside phagolysosomes. **(A)** dPLB cells were stained with LAMP-1 (red) after infection with wild-type or Δ*pksP c*onidia (FITC labeled; green) for 4 h. Images are representative of three biological replicates. **(B)** Quantification of LAMP-1 co-localization with CFW-labelled conidia in phagolysosomes and **(C)** Percentage of acidified conidia in dPLB cells after 2- and 4-hours post-infection with wild-type and Δ*pksP c*onidia. Data are presented as mean ±SD; *** = p < 0.001. **(D)** Colocalization of conidia (CFW; blue) in acidified compartments labeled with Lysotracker (red). Images are representative of three biological replicates. Scale bars are 10 µm.

The interference of the acidification pathway in alveolar macrophages by fungal DHN-melanin occurs through the inhibition of V-ATPase assembly (26, 27). This multiprotein complex plays a major role in lowering the pH from 6 to <4.5 by pumping H^+^ ions across the phagolysosomal membrane. Staining of the V-ATPase V1 subunit in dPLB cells revealed that the percentage of recruitment to conidia-containing phagolysosomes was fairly similar for wild-type and *ΔpksP* conidia (**Fig. S3A to B**), suggesting that other proton pumps may contribute to acidification of these phagolysosomes in neutrophil-like cells. An additional defense mechanism for intra-phagosomal degradation of pathogens occurs via reactive oxygen species (ROS) production. As expected, we observed dPLB cells to be capable of producing ROS after staining with CellROX Orange (**Fig. S3C**), consistent with the literature (28, 29).

### *A. fumigatus* triggers formation of NETs from dPLB cells

One defense mechanism employed by neutrophilic granulocytes to fight against pathogens is the formation of NETs. These structures consist of condensed chromatin and various enzymes that are released into the extracellular space and typically correlate with death of the cell (30). NETs contain several anti-fungal proteins, which are responsible for the fungistatic effect against *A. fumigatus* hyphae (15). To test if dPLBs are also able to form NETs in response to *A. fumigatus* and serve as a model for this pathway, we co-incubated dPLBs cells with hyphae and stained for nucleic acid with 4′,6-diamidino-2-phenylindole (DAPI; **Fig. 3A**). As a positive control, we induced NET formation using phorbol myristate acetate, a known trigger of NET formation in PLB-985 cells (**Fig. 3B**; (21, 31)). We observed the presence of histone H3 embedded in the DNA fibers and an association of NETs with fungal hyphae (**Fig. 3B**). Taken together these results showed that *A. fumigatus* is able to trigger NET formation in dPLB cells and that these NETs are specifically directed against the hyphae.

**FIG 3.**
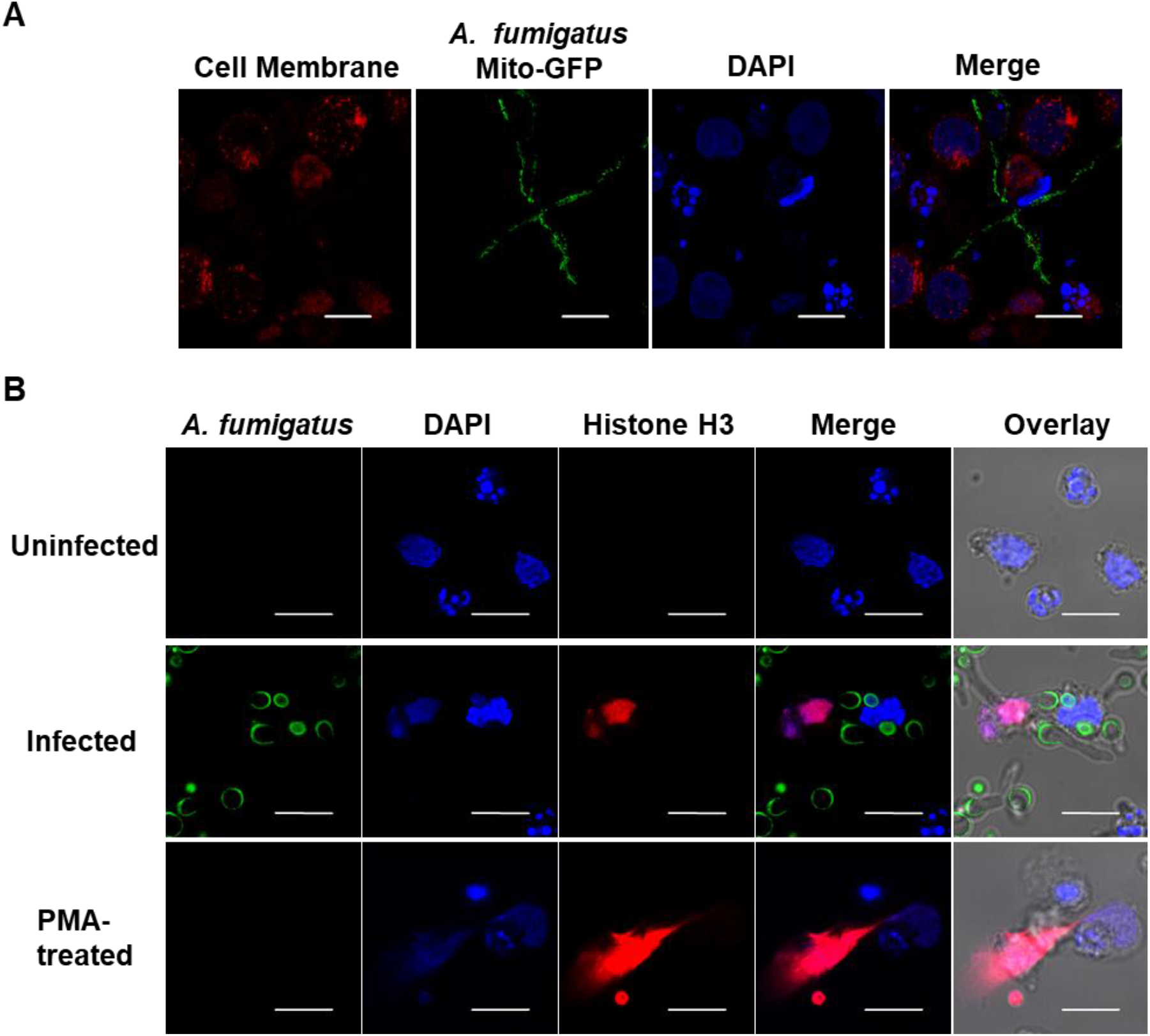
Production of NETs in response to *A. fumigatus* infection. **(A)** Confocal scanning laser microscopy of nucleic acid stained with DAPI (blue) released from dPLB cells stained with Cell Mask (red) after challenge with fungal hyphae containing a mitochondrial GFP reporter (*A. fumigatus* strain AfS35 / pJW103 expressing a mitochondrial GFP reporter; green). Phorbol myristate acetate (PMA) was used as a positive control. Data representative of three biological replicates. **(B)** Confocal micrographs of NET markers, Histone H3 (red) and nucleic acid stained with DAPI (blue), produced by dPLB cells after contact with *A. fumigatus* hyphae (FITC-labeled; green). Images are representative of three biological replicates. Scale bars are 10 µm.

### dPLB cells produce extracellular vesicles in response to infection

Recently, a new defense mechanism from neutrophilic granulocytes was discovered, the production of antimicrobial extracellular vesicles (32, 33). These small lipid-enclosed nanoparticles are released from primary PMNs after contact with microorganisms and, in the case of *A. fumigatus* conidia, can inhibit fungal growth after coincubation (10). This effect is likely due in part to their protein cargo, which consists of antimicrobial peptides such as myeloperoxidase, azurocidin, and cathepsin G. To assess if dPLB cells can produce extracellular vesicles spontaneously or in response to *A. fumigatus*, we incubated cells with or without conidia for 2 and 4 hours. Next, we isolated extracellular vesicles using two different methods; first using a previously described differential centrifugation-based approach (DC; (10)) that enriches for medium-sized extracellular vesicles and a second approach that relies on size-exclusion chromatography (SEC) to purify a more selective population of smaller extracellular vesicles. Using these methods, dPLB cells were observed to actively secrete extracellular vesicles over time in a manner comparable to primary human PMNs (**Fig. 4A to B)**. Using nanoparticle tracking analysis, we observed the median size of particles to be around 200 nm, similar to extracellular vesicles derived from primary neutrophils, a feature that was comparable between infection-derived and spontaneously released extracellular vesicles (**Fig. 4C to D**).

**FIG 4.**
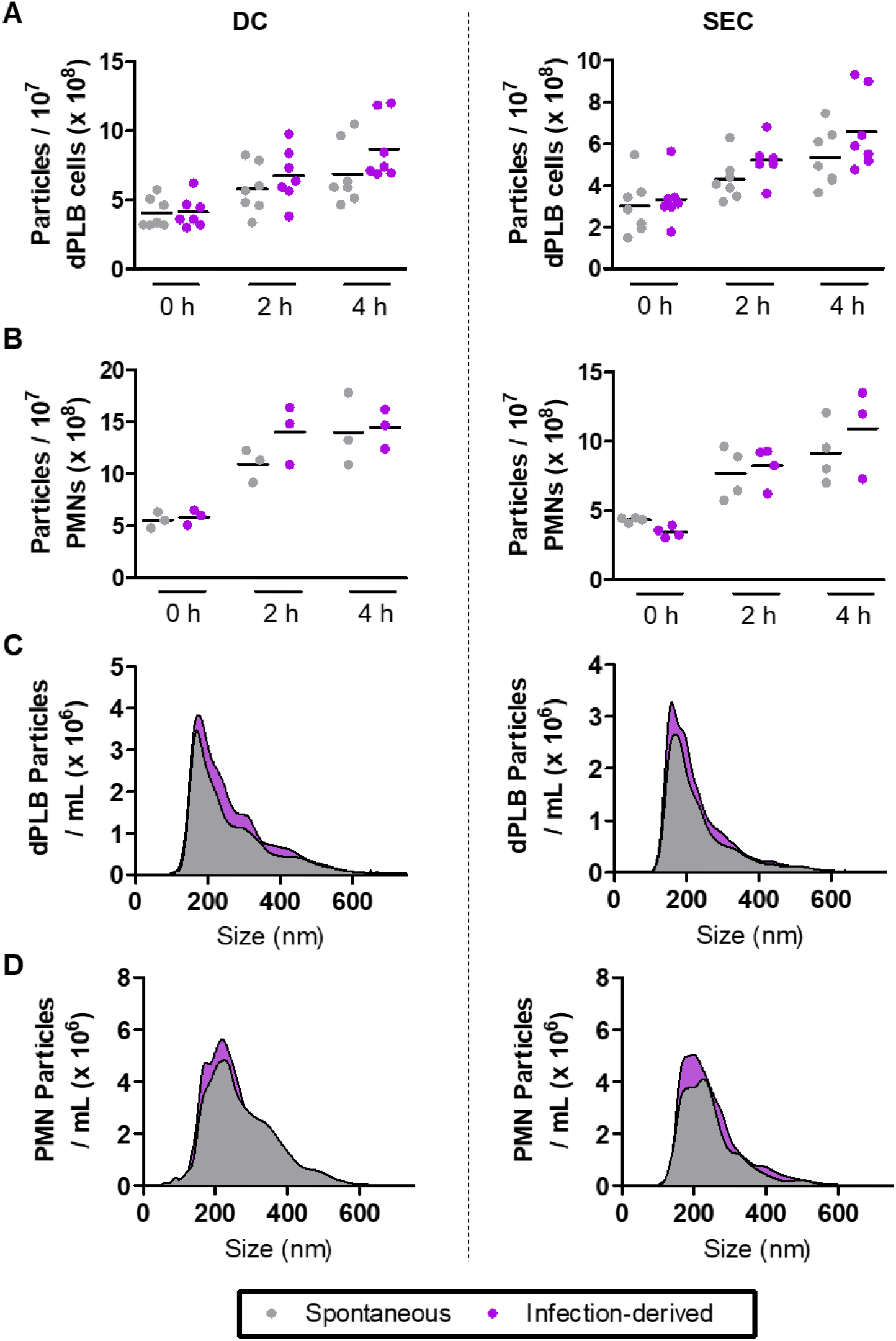
dPLB cells produce extracellular vesicles comparable in size to primary human PMNs. Extracellular vesicles released spontaneously or produced in response to infection with opsonized *A. fumigatus* conidia were isolated using a differential centrifugation-based approach (DC) or a size-exclusion chromatography-based approach (SEC) and quantified using nanoparticle tracking analysis. Extracellular vesicles were quantified from **(A)** dPLB cells or **(B)** primary human PMNs at 0, 2, and 4 hours post infection and show the average of at least six biological replicates and three biological replicates, respectively. Representative size histograms from five biological replicates are shown for extracellular vesicles derived from **(C)** dPLB cells and **(D)** primary human PMNs.

We next compared the protein content of dPLB-derived extracellular vesicles isolated, at the 2 hours timepoint, using each isolation method with and without fungal infection by LC-MS/MS-based proteomics analysis. We were able to identify 1,984 unique proteins across all four samples (**Dataset S1**). The majority of identified proteins (737 proteins) were found in all four samples (**Fig. 5A**). We expected that isolation using size-exclusion chromatography would improve the quality of the isolated particles as has been shown previously (34), and this was in fact the case. We observed an increase in extracellular vesicle markers like the tetraspanins CD63 and CD81, and tumor susceptibility gene 101 (TSG101), and a decrease in cytoplasmic proteins like calnexin (CANX; **Table 1**).

**Table 1.** RMN-fold change of selected proteins identified in extracellular vesicles (EVs) from dPLB cells. Abbreviations: DC, differential centrifugation-based approach; SEC, size-exclusion chromatography-based approach.

| Gene | Accession | Infection-derived EVs (DC) / Spontaneous EVs (DC) | Infection-derived EVs (SEC) / Spontaneous EVs (SEC) | Infection-derived EVs (SEC) / Infection-derived EV (DCs) | Spontaneous EVs (SEC) / Spontaneous EVs (DC) |
| --- | --- | --- | --- | --- | --- |
| *EV Markers* |  |  |  |  |  |
| **CD63** | F8VWK8 | 1.274 | 1.424 | 1.478 | 1.323 |
| **TSG101** | F5H442 | 1.215 | 1.630 | 1.554 | 1.158 |
| **CD81** | E9PRJ8 | 1.150 | 1.413 | 2.055 | 1.673 |
| *Non-EV Markers* |  |  |  |  |  |
| **CANX** (Calnexin) | P27824 | -1.031 | -1.082 | -1.468 | -1.398 |
| **MPO**<br>(Myeloperoxidase) | P05164 | -1.024 | 1.026 | 2.063 | 1.963 |
| **CTSG**<br>(Cathepsin G) | P08311 | 1.045 | 1.042 | 1.117 | 1.120 |
| **AZU1**<br>(Azurocidin) | P20160 | -1.141 | 1.029 | 1.521 | 1.295 |

**FIG 5.**
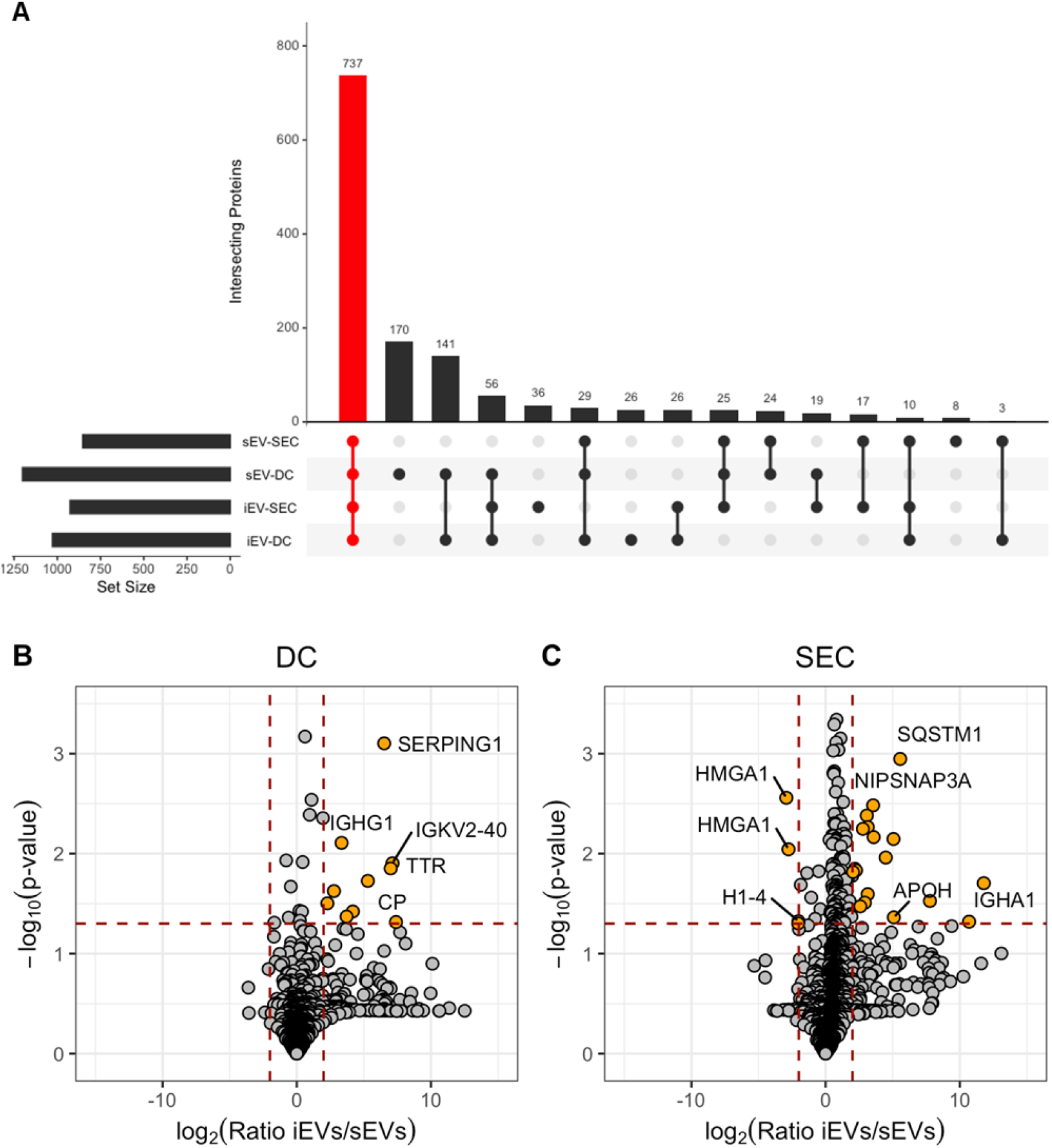
SEC enriches for extracellular vesicle populations. LC-MS/MS proteomics analysis was performed on extracellular vesicles isolated from dPLB cells using a differential centrifugation-based approach (DC) or a size-exclusion chromatography-based approach (SEC) in the presence or absence of infection with opsonized *A. fumigatus* conidia. **(A)** Proteins identified in spontaneously released extracellular vesicles (sEVs) and infection-derived extracellular vesicles (iEVs) from at least two replicates of a given sample were intersected using UpSetR. The red bar indicates proteins that were found in all four samples. Volcano plots show the log_2_ ratio of infection-derived extracellular vesicles (iEVs) versus spontaneously released EVs (sEVs) for **(B)** DC-based isolation and **(C)** SEC-based isolation. Input data included values from all replicates using the RMN data included in **Dataset S1**. Plots were created using ggplot2 in R. Proteomics data is from three analytical replicates of three independent biological replicates. Orange circles represent proteins with greater than 2-fold change and p = < 0.05. Selected proteins are named for clarity.

Extracellular vesicles produced by primary PMNs in response to *A. fumigatus* infection are known to be enriched for antimicrobial cargo proteins like azurocidin, cathepsin G, and defensin (10). In line, with the data from primary neutrophils we found that dPLBs also contained many of the same proteins in both infection-derived and spontaneously released extracellular vesicles (**Table 1**), as evidenced by an UpSetR plot showing overlapping proteins cohorts (**Fig. S4;** (10)), although with some exceptions. For example, dPLBs appeared to lack defensin, neutrophil elastase, and some histone proteins previously observed. Ultimately, improvements in LC-MS/MS technology and the advantage of higher input amounts of extracellular vesicle protein using large amounts of dPLB cells in culture resulted in significantly more proteins detected in the dataset provided here compared to efforts using primary neutrophils (10), proving that dPLBs offer a scalable system for the elucidation of novel mechanisms of neutrophil extracellular vesicle biology.

### dPLB extracellular vesicles produced against *A. fumigatus* limit fungal growth

The most compelling feature of infection-derived extracellular vesicles of human neutrophils is likely their antifungal capacity, as we previously reported (10). We set out to determine if the extracellular vesicles produced by dPLB cells in response to *A. fumigatus* opsonized conidia are antifungal to a mitochondrial-GFP reporter strain used previously as a marker of fungal viability (35). First, we incubated the reporter strain with dPLB cells and primary human PMNs and assessed the capacity of the cells to control infection. PMNs were capable of minimizing fungal outgrowth after 22 hours, whereas dPLBs did not completely contain fungal growth (**Fig. 6A**). We then assessed the antifungal capacity of the dPLB extracellular vesicles. For this experiment, extracellular vesicles from equal numbers of dPLBs and primary human PMNs were isolated using differential centrifugation and found to be antifungal against the mitochondrial-GFP reporter strain. This was evidenced by fragmentation of fungal mitochondria after administration of extracellular vesicles to germinating conidia, which were allowed to germinate for 6 hours prior to overnight incubation with extracellular vesicles, 3 mM H_2_O_2_ as a positive control, or left untreated as a negative control (**Fig. 6B**). Although, the mitochondrial reporter clearly indicated fragmentation, the extracellular vesicles from dPLB did appear to be in some cases less effective in limiting the length development of the hyphae than those from primary PMNs. It is important to note that dPLBs produced slightly fewer extracellular vesicles, which could potentially explain the decreased antifungal activity in these assays (**Fig. 6B**). Additional representative images of dPLB-derived extracellular vesicles are shown to indicate the spectrum of phenotypes observed for each experimental condition (**Fig. S5**). Collectively, these results suggest that dPLB infection-derived extracellular vesicles can limit *A. fumigatus* hyphal growth, similar in fashion to primary human PMNs (10).

**FIG 6.**
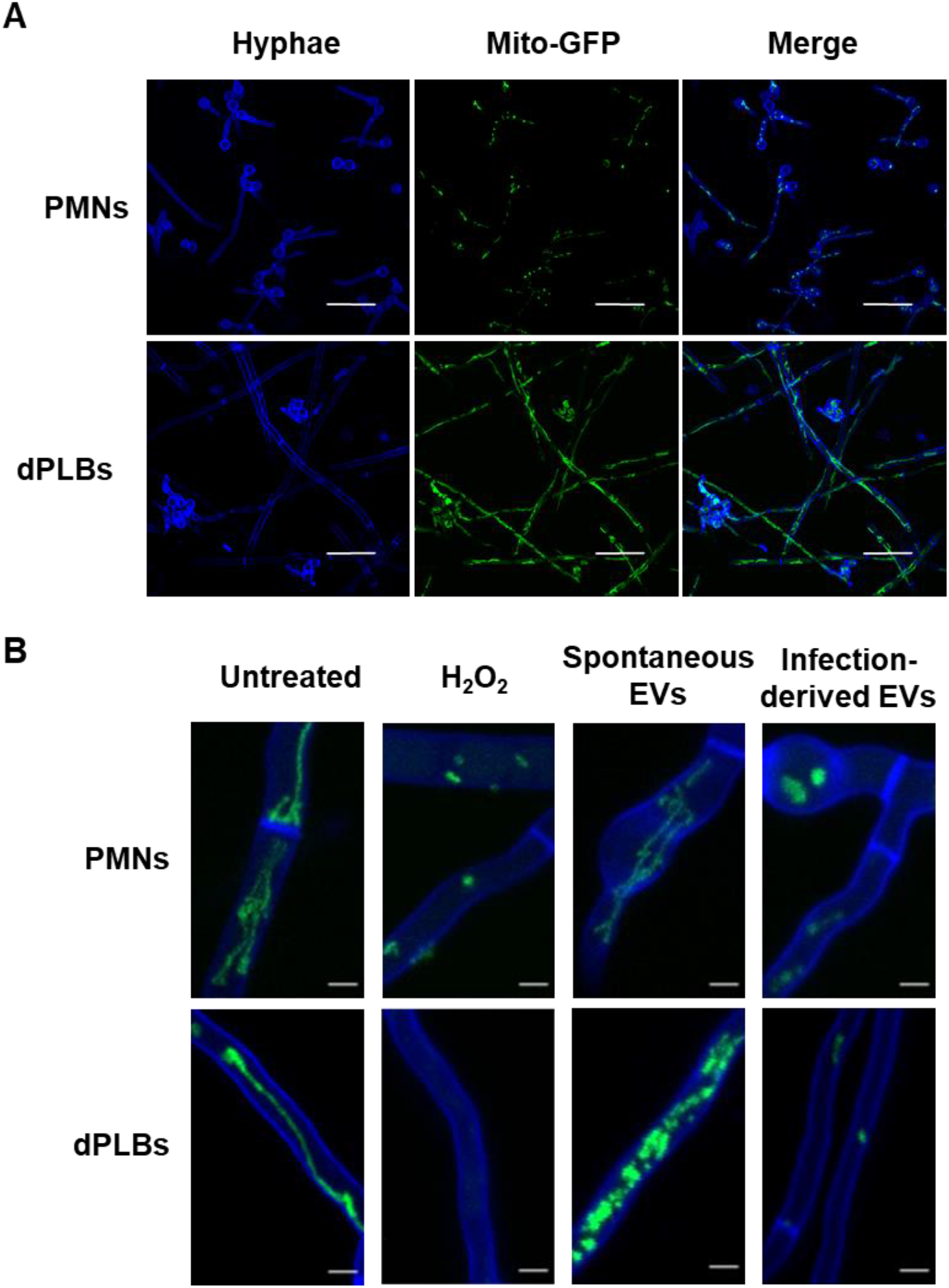
Infection-derived extracellular vesicles from dPLBs are antifungal to *A. fumigatus* hyphae. **(A)** Conidia from *A. fumigatus* strain AfS35 containing plasmid pJW103 expressing a mitochondrial GFP reporter (green) were opsonized and co-incubated with freshly harvested human PMNs or dPLBs. After overnight incubation (22 hours) samples were stained with CFW for 10 min and images were taken using a Zeiss LSM 780 confocal microscope. Scale bars are 20 µm. **(B)** The *A. fumigatus* strain AfS35 expressing a mitochondrial GFP reporter (green) was first grown for 6 hours and then stained with calcofluor white (blue) and incubated overnight with spontaneously released extracellular vesicles or infection-derived extracellular vesicles isolated from primary human PMNs or dPLBs. As a control, untreated hyphae and hyphae treated with 3 mM H_2_O_2_ to induce cell death are included. An intact mitochondrial network is shown by a filamentous network, whereas a disrupted network is shown by fragmentation or the lack of green signal. Scale bars are 5 µm.

## MATERIALS AND METHODS

### Fungal strains cultivation and opsonization

Cultivation of *A. fumigatus* strains CEA10 (Fungal Genetics Stock Center; A1163), CEA17 Δ*akuB*^*ku80*^ Δ*pksP* (36), and AfS35/pJW103 (35) was performed on malt agar plates (Sigma-Aldrich) supplemented to a final concentration of 2% (wt/vol) agar for 5 days at 37 °C. Conidia were harvested in sterile, ultrafiltrated water, filtered through a 30-µm pore filter (MACS, Miltenyi Biotec). Prior to confrontation with PLB-985 cells, conidia were opsonized with normal human serum (Merck Millipore). Briefly, 900 µl of spore suspension was mixed with 100 µl of normal human serum and incubated in a thermomixer at 37 °C for 30 min shaking at 500 rpm. The spore suspension was washed three times by collecting spores *via* centrifugation at 1800 x *g* for 1 min at 4 °C and resuspending in fresh PBS 1X. Following washing, spores were enumerated in a Thoma chamber in preparation for infection assays.

### Cell culture cultivation, differentiation, and infection

PLB-985 cells were maintained in Roswell Park Memorial Institute medium 1640 (RPMI; Gibco) supplemented with 10% (vol/vol) fetal bovine serum (FBS; HyClone, GE Life science), 100 U/ml penicillin-streptomycin antibiotic solution (Lonza) and 2 mM UltraGlutamine (alanyl L-glutamine; Gibco). For differentiation into a neutrophilic granulocyte cell type, 4×10^5^ cells/ml were resuspended in supplemented RPMI medium containing 2.5% (vol/vol) FBS and 0.5% DMF (Sigma-Aldrich) for 4 days at 37 °C with 5% (vol/vol) CO_2_. On day 4, fresh differentiation medium was added, and the cells were incubated for 3 additional days. On day 7, cells were collected, centrifuged at 300 x *g* for 5 min and then seeded in wells for experiments at the appropriate concentrations. The medium used for all the assays was composed of RPMI supplemented with 1% (vol/vol) exosome-depleted FBS (Life Technologies GmbH), 2 mM UltraGlutamine, and 0.5% (vol/vol) DMF.

### Isolation of human PMNs

Fresh venous blood was drawn from adult healthy volunteers, aged 20 to 35 years, after obtaining written consent. This study was approved by the Jena University Hospital Institutional Review Board (approval number 5074-02/17) in agreement with the Declaration of Helsinki. Blood was collected in EDTA BD Vacutainer tubes (BD Biosciences) and mixed with PBS 1X. 6 ml of this solution was carefully layered onto (avoiding any mixing) 5 ml (1:1 vol/vol) of PolymorphPrep solution (Progen) and centrifuged 50 min at 500 x *g* for gradient separation. After centrifugation the neutrophil layer was collected, mixed with an equal volume of PBS 0.5% and re-centrifuged for 15 min at 400 x *g*. To limit contamination of red blood cells, the pellet was lysed with the ACK lysis buffer (Gibco) and centrifuged as before. The PMNs were finally re-suspended in RPMI medium without phenol red (Thermo Fischer scientific) and after addition of Trypan blue counted with a Luna automated cell counter (Logos biosystem). For each experiment the viability of neutrophils was ≥ 95%.

### Determination of cell viability

To assess cell viability after confrontation with *A. fumigatus* conidia, the release of LDH was determined using the CyQUANT™ LDH cytotoxicity assay kit (Thermo Fisher Scientific) according to the manufacturer’s instruction. Cells treated with *A. fumigatus* conidia with a multiplicity of infection (MOI) of 5 for 2 hours were tested for their LDH activity, which was compared to the spontaneous LDH activity and the maximum LDH activity. Release of LDH was below the limit of detection for all three independent biological replicates performed.

### Phagocytosis assays

To assess phagocytic ability, we used a combination of imaging flow cytometry and confocal fluorescence microscopy. For both methods the conidia were first stained with fluorescein isothiocyanate (FITC) and then after confrontation of dPLBs or primary neutrophils, counterstained with calcofluor white (CFW; Sigma -Aldrich). The FITC solution was obtained by dissolving FITC powder (Sigma-Aldrich) in 5 mL of 0.1 M sodium carbonate (Na_2_CO_3_) followed by filtration through a 0.22-µm pore size filter (Carl Roth). Afterwards 1 ml of this solution was mixed with 500 µl of spore suspension and incubated for 30 min at 37 °C while shaking at 1000 rpm in the dark. The spores were then pelleted and washed three times with PBS 1X with 0.001% (vol/vol) Tween 20 (Carl Roth). During the last washing step Tween was removed to avoid residual detergent in the samples (25, 37). Before counting, the conidia were opsonized following the protocol described above. They were then added to the cells and co-incubated for 2 and 4 hours at 37 °C with 5% (vol/vol) CO_2._ At the end of each time point CFW was added to a final concentration of 1 µg/ml and incubated for 1 min. Afterwards the cell suspension was transferred to microcentrifuge tubes, centrifuged at 600 x *g* for 2 min at 4 °C and washed with PBS 1X twice. After discarding the supernatant, the pellet was fixed with 150 µL of 3.7% (vol/vol) formaldehyde in PBS at room temperature for 15 min and subsequently washed again as described above. For imaging flow cytometry measurements, the cells were resuspended in 150 µL of PBS 1X and analyzed immediately or stored at 4 °C for no more than 24 hours. Four independent experiments were performed and for each replicate 5000 cells were analyzed using the ImageStream X Mark II (Luminex). A 488 nm laser was used to detect FITC staining and a 405 nm laser for the CFW. The laser voltage was adjusted depending on the sample. For wild-type conidia, we used 10 mW of 488 nm laser and 2 mW for the 405 nm laser, and for Δ*pksP* conidia 1 and 2 mW of the respective lasers. The fluorescence intensities of the samples were compensated and analyzed with the IDEAS software (Luminex). An example of the gating strategy used for analysis can be found in the supplement files (**Fig. S1B to C**). For the fluorescence microscopy analysis, cells were seeded in 8-well µ-slides (Ibidi) and visualized using a Zeiss LSM 780 confocal microscope (Carl Zeiss) from at least three biological replicates as described above.

### Immunofluorescence assays

For immunofluorescence, cells were allowed to adhere in a 24-well plate with poly-L-lysine-coated glass coverslips for 1 hour (Merck). After infection with labelled *A. fumigatus* spores and incubation for the noted times, cells were fixed using 3.7% (vol/vol) formaldehyde in PBS for 10 min, rinsed three times with PBS 1X, permeabilized for 15 min using 0.1% (vol/vol) Triton X-100 in PBS 1X or 0.1% (wt/vol) saponin in PBS 1X, and then blocked for 30 min with 2% (wt/vol) bovine serum albumin (BSA). After permeabilization, washed cells were incubated with primary rabbit anti-Lamp-1 antibody (Abcam 24170, 1:100 dilution), anti-V-ATPase V1 subunit antibody (Abcam 73404, 1:100 dilution), or anti-Histone H3 (Cell signaling DIH2, 1:200 dilution) antibody in 1% (wt/vol) BSA in PBS 1X, followed by incubation with secondary goat anti-rabbit IgG antibody DyLight 633 (Thermo Fisher Scientific). The glass cover slips were mounted onto microscopy slides and visualized using a Zeiss LSM 780 confocal microscope (Carl Zeiss). LAMP-1 recruitment was quantified by comparing the positive signal from stained phagolysosomes to non-stained phagolysosomes.

### Reactive oxygen species measurements

ROS production was detected in dPLB cells by seeding cells in 8-well µ-slides (Ibidi) coated with poly-L-lysine followed by infection with *A. fumigatus conidia* at an MOI of 5 for either 2 or 4 hours. CellROX® Orange Reagent (Thermo Fisher Scientific) was added 30 minutes prior the end of the infection time points, and ROS production was imaged using a Zeiss LSM 780 confocal microscope (Carl-Zeiss) for fluorescent intensity.

### Cytokine measurement

For the detection of IL-8 and IL-1β, 2×10^5^ dPLB cells or primary neutrophils were seeded in 24-well plates. After addition of conidia at a MOI of 5, plates were incubated at 37 °C with 5% (vol/vol) CO_2_ for 2, 4, or 6 hours. An additional 24-hour timepoint was included for dPLB cells. As a positive control, cells were treated with 5 µg/mL of lipopolysaccharide (LPS; Sigma-Aldrich L4516). At the appropriate end point, samples were collected, centrifuged at 300 x *g* for 5 min to remove cell debris, and frozen at -20°C and analyzed within 3 days. Cytokines were measured using human ELISA Max deluxe kits (BioLegend) according to the manufacturer’s instruction.

### Acidification assays

Acidification assays were performed as mentioned previously (27). Briefly, the cells were incubated with 50 nM lysotracker DND-99 (Thermo Fisher Scientific) for 1 hour prior to infection. After that, FITC-labeled conidia were added to the cells for 2 or 4 hours and imaged using Zeiss LSM 780 confocal microscope (Carl Zeiss). For quantification, 200 conidia-containing phagolysosomes were counted and evaluated for acidification. The values represent the mean ± SD of three separate experiments.

### Extracellular vesicle isolation

After infection with *A. fumigatus* the supernatant of dPLB cells was collected, and extracellular vesicles were isolated using two different methods: a differential centrifugation-based approach or a size exclusion chromatography-based approach. The first method was described previously for the isolation of neutrophil-derived extracellular vesicles (10, 32). In both approaches samples were centrifuged at 3000 x *g* for 15 min at 4 °C and then filtered through 5-µm pore size filters (Carl Roth). In the first method, samples were then centrifuged at 19,500 x *g* for 20 min at 4 °C to collect extracellular vesicles. For the second approach, the clarified filtrate was concentrated using Amicon Ultra-15 centrifugal filters (Merck) with a molecular mass cut-off (MWCO) of 100 kDa for 10 min at 4 °C and 3220 *x g* and finally loaded on size-exclusion chromatography qEV 70 nm columns (Izon). After discarding the 3 mL void volume, 1.5 mL of extracellular vesicle sample were collected and measured. When necessary, extracellular vesicles were further concentrated using 10 kDa cutoff Amicon Ultra-0.5 ml filters (Merck). Extracellular vesicles were used fresh for most of the downstream experiments except for proteomic analysis where they were frozen at -20 °C.

### Nanoparticle tracking analysis

Particle concentration and size distribution were analyzed using a nanoparticle tracking analysis (NTA) NS300 device with a 650 nm laser (Malvern Instruments Ltd). Fresh samples were measured with a constant flow rate of 20 and a temperature set to 25 °C. Five 60-second videos were recorded with a camera level of 11. Videos were then analyzed with the NTA 3.2.16 software using a detection threshold of 4.

### Protein preparation for LC-MS/MS

After isolation as described above, extracellular vesicles were delipidated as described previously (10) using the protein precipitation protocol of Wessel and Flügge (38). Delipidated extracellular vesicles were resolubilized in 100 µl of 50 mM triethyl ammonium bicarbonate (TEAB) in 1:1 (vol/vol) trifluoroethanol/water. For reduction of cysteine thiols, the solution was mixed with 10 mM Tris(2-carboxyethyl) phosphine and alkylated with 12.5 mM chloroacetamide at 70 °C for 30 min in the dark. Proteins were digested for 18 hours at 37 °C with trypsin/LysC mix (Promega) at a protein-to-protease ratio of 25:1. Tryptic peptides were first completely evaporated using a vacuum concentrator (Eppendorf) and then resolubilized in 0.05% (vol/vol) trifluoroacetic acid (TFA) in 2:98 (vol/vol) acetonitrile/water. Finally, the samples were filtered through Ultrafree-MC Hydrophilic PTFE membrane (0.2-µm pore size) spin filters (Millipore) and stored at -20 °C until measurement. Each sample was measured in triplicate (3 analytical replicates of 3 biological replicates) as follows.

### LC-MS/MS analysis

LC-MS/MS analysis was performed on an Ultimate 3000 nano RSLC system connected to a QExactive HF mass spectrometer (both Thermo Fisher Scientific). Peptide trapping for 5 min on an Acclaim Pep Map 100 column (2 cm x 75 µm, 3 µm) at 5 µL/min was followed by separation on an analytical Acclaim Pep Map RSLC nano column (50 cm x 75 µm, 2µm). Mobile phase gradient elution of eluent A (0.1% (vol/vol) formic acid in water) mixed with eluent B (0.1% (vol/vol) formic acid in 90/10 acetonitrile/water) was performed as follows: 0 min at 4% B, 30 min at 12% B, 75 min at 30% B, 85 min at 50% B, 90-95 min at 96% B, 95.1-120 min at 4% B.

Positively charged ions were generated at spray voltage of 2.2 kV using a stainless-steel emitter attached to the Nanospray Flex Ion Source (Thermo Fisher Scientific). The quadrupole/orbitrap instrument was operated in Full MS / data dependent MS2 (Top15) mode. Precursor ions were monitored at m/z 300-1500 at a resolution of 120,000 FWHM (full width at half maximum) using a maximum injection time (ITmax) of 120 ms and an AGC (automatic gain control) target of 3×10^6^. Precursor ions with a charge state of z=2-5 were filtered at an isolation width of m/z 1.6 amu for further HCD fragmentation at 27% normalized collision energy (NCE). MS2 ions were scanned at 15,000 FWHM (ITmax=100 ms, AGC= 2×10^5^). Dynamic exclusion of precursor ions was set to 25 s and the underfill ratio was set to 1.0%. The LC-MS/MS instrument was controlled by Chromeleon 7.2, QExactive HF Tune 2.8 and Xcalibur 4.0 software.

### Database search and data analysis

Tandem mass spectra were searched against the UniProt database of *Homo sapiens* (https://www.uniprot.org/proteomes/UP000005640; 2021/05/17) and *Neosartorya fumigata* (https://www.uniprot.org/proteomes/UP000002530; 2021/05/17) using Proteome Discoverer (PD) 2.4 (Thermo) and the algorithms of Mascot 2.4.1 (Matrix Science), Sequest HT (version of PD2.2), MS Amanda 2.0, and MS Fragger 3.2. Two missed cleavages were allowed for the tryptic digestion. The precursor mass tolerance was set to 10 ppm and the fragment mass tolerance was set to 0.02 Da. Modifications were defined as dynamic Met oxidation, and protein N-term acetylation with and without methionine-loss as well as static Cys carbamidomethylation. A strict false discovery rate (FDR) < 1% (peptide and protein level) were required for positive protein hits. If only 1 peptide per protein has been identified the hit was accepted if the Mascot score was >30 or the MS Amanda score was >300 or the Sequest score was >4 or the MS Fragger score was >8. The Percolator node of PD2.4 and a reverse decoy database was used for qvalue validation of spectral matches. Only rank 1 proteins and peptides of the top scored proteins were counted. Label-free protein quantification was based on the Minora algorithm of PD2.2 using a signal-to-noise ratio >5. Imputation of missing quan values was applied by setting the abundance to 75% of the lowest abundance identified for each sample. Normalization was based on a replicate median total peptide sum approach, which was calculated based on the sum of all identified peptide abundance values per replicate sample. The sums of each of the three replicates from the four sample groups were used to calculate median values. Normalization factors were calculated by dividing median values of the respective sample group by the abundance sum of each sample. Normalization factors were multiplied with single protein abundance values of each replicate/sample. The p-values are based on a Student’s t-test. Ratio-adjusted p-values were calculated by dividing p-values with the log4ratio of the protein abundance levels. Significant differences in protein abundance were defined when the following three requirements were reached: At least a 4-fold change in abundance (up and down), a ratio-adjusted p-value <0.05, and at least identified in 2 of 3 replicates of the sample group with the highest abundance. Intersection plots were created using the UpSetR package (39) and only include proteins that were detected in at least two replicates of a given sample. Volcano plots were created using ggplot2 in R using the replicate median total peptide sum normalized (RMN) data for all proteins detected in **Dataset S1**.

### Fungal mitochondrial reporter inhibition assay

To characterize growth inhibition and killing of hyphae by fungal-induced extracellular vesicles an *A. fumigatus* strain expressing a mitochondrial GFP reporter (AfS35/pJW103; (35)) has been used as described previously (10). Extracellular vesicles produced by 1×10^7^ dPLB cells or PMNs were isolated and co-incubated with the reporter strain. After 16 hours of incubation samples were stained with CFW and images were taken using Zeiss LSM 780 confocal microscope (Carl Zeiss).

### Data availability

The mass spectrometry proteomics data have been deposited to the ProteomeXchange Consortium via the PRIDE partner repository (40) with the dataset identifier PXD027032.

### Statistical Analysis

Data were plotted and statistically analyzed using GraphPad Prism software 5.0 (GraphPad Software) unless otherwise noted. The Student’s t test was used for significance testing when comparing two groups. Differences between the groups were considered significant at a P value of <0.05. Throughout the article, significance is denoted as follows: *, P = <0.05; **, P = <0.01, ***, P = <0.001; ns, nonsignificant.

### Ethics Statement

Peripheral human blood was collected from healthy volunteers only after written informed consent was provided. The study was conducted in accordance with the Declaration of Helsinki and approved by the Ethics Committee of the University Hospital Jena (approval number 5074-02/17).

## DISCUSSION

Neutrophils are critical players in the immune response to fungal pathogens stemming from their numerous antimicrobial capacities (10, 41). To study neutrophil functions, numerous cell lines that model these abilities *in vitro* have been considered with limited success. The most promising remains the HL-60 leukemia cell line and PLB-985 sublineage, which have been used as models for aspergillosis and bacterial killing (21-23, 28). In the present study, we analyze different aspects of the interaction between DMF-differentiated PLB-985 myeloid cells and the fungus *A. fumigatus* for the first time, making frequent comparisons with data from primary neutrophils. Primary neutrophils are capable of phagocytosing and processing *A. fumigatus* conidia, features that were maintained in dPLB cells. As expected for cultured cells, the observed phagocytosis rate of *A. fumigatus* conidia by dPLB cells was lower than that of primary human PMNs. It is possible that additional differentiation methods will ultimately reveal higher rates of phagocytosis, as PLB-985 cells differentiated with DMF are known to exhibit lower phagocytic activity than cells differentiated with dimethyl sulfoxide (42). In terms of cytokine release, we observed increased release of IL-8 from dPLBs after infection with *A. fumigatus* opsonized conidia, but not IL-1β, consistent with our observations in human PMNs. These results are also in agreement with those collected from primary PMNs co-incubated with L-ficolin opsonized conidia for 8 hours, where secretion of IL-8 was also much higher than IL-1β (43).

We next stained dPLB cells for the endosomal marker LAMP-1 and measured the acidification of the phagolysosomes via Lysotracker to assess intracellular processing of fungal conidia. These experiments confirmed that around 20% of wild-type conidia are completely internalized inside the phagosomes and that lysosomal fusion occurs as a next step of conidial processing. Interestingly, and as already demonstrated for primary monocytes and neutrophils (25), melanized conidia showed less acidification around the phagolysosomal membrane than the mutant lacking DHN-melanin. This effect in monocytes is caused by inhibition of V-ATPase assembly. Surprisingly, staining for the V1 subunit of the V-ATPase did not reveal a significant difference between the wild-type and Δ*pksP* strains in dPLB cells as has been observed in other cell systems. These results suggest that neutrophils potentially employ additional mechanisms to acidify phagolysosomes in neutrophilic granulocytes after *A. fumigatus* infection or that the methods used were not sufficiently sensitive to detect slight differences in acidification. Together, these results suggest that dPLBs can serve as an intriguing model to study various aspects of neutrophil phagocytosis and intracellular processing in a more tractable *in vitro* model system.

A conidium that escapes phagocytosis can sometimes germinate into a hypha, a morphotype that is much more difficult to eliminate and requires the antifungal activity of neutrophils. One mechanism that aids in control of *A. fumigatus* hyphae is the release of fungistatic NETs composed of DNA and antimicrobial proteins. By staining for extracellular DNA and NET constituent histone H3, we could clearly show that dPLB cells also produce NETs in response to *A. fumigatus* hyphae, comparable to primary human PMNs and in agreement with other studies challenging dPLBs with bacterial pathogens (21, 22).

Production of extracellular vesicles from primary neutrophils co-incubated with *Aspergillus* conidia or bacteria like *Staphylococcus aureus* is an important defense strategy against microorganisms (10, 32). We found that dPLB cells were also able to generate comparable populations of extracellular vesicles to PMNs upon contact with opsonized conidia. Accumulation of extracellular vesicles of approximately 200 nm in diameter was observed over time, independently of the isolation method we applied. Proteomic analysis revealed that by using a size-exclusion chromatography-based isolation approach, the extracellular vesicles population could be enriched for extracellular vesicle marker proteins like CD63, CD81, and TSG101. We observed that the samples obtained by this method had a higher abundance of proteins and showed more differences between the spontaneously released and infection-derived extracellular vesicles. In particular, infection-derived extracellular vesicles were enriched in immunoglobulins and complement factors when compared to spontaneously released vesicles. These serum proteins may be introduced to the infection system during opsonization of the fungal conidia despite thorough washing. It remains unclear whether these immunoglobulins and complement factors play an important role in extracellular vesicle biology during *A. fumigatus* infection. As a hint for such a role, the association of immunoglobulins and complement proteins was shown previously to be important in systemic lupus erythematosus (SLE) disease, where extracellular vesicles with immunoglobulin cargo could act as immune complexes to mediate inflammation (44, 45).

In the proteomic analysis of extracellular vesicles derived from primary neutrophils infected with *A. fumigatus*, several antimicrobial peptides, such as cathepsin G and azurocidin, were detected (10). Co-incubation of these extracellular vesicles from primary neutrophils with fungal hyphae arrested growth, which was confirmed in this study. Using dPLB cells, we identified extracellular vesicles containing cathepsin G and azurocidin; however, the distribution remained relatively equal between spontaneously released and infection-derived extracellular vesicles. We suspect that this is due to the differentiation procedure of dPLB cells with DMF, which provides a mild inflammatory stimulus. Nevertheless, we observed mitochondrial damage in hyphae treated with infection-derived but not spontaneously released extracellular vesicles. We can postulate several reasons for this result. First, it is possible that immunoglobulins are playing a role in targeting extracellular vesicles to fungal hyphae. In this case, the protein cargo may be the same, but targeting would be inefficient in spontaneously released extracellular vesicles. Of note, opsonization of the bacterial pathogen *Staphylococcus aureus* was not found to influence the antimicrobial capacity of neutrophil-derived extracellular vesicles (32). A second option is that additional extracellular vesicle cargo molecules such as RNA or lipids may play a role in the antifungal activity. It has been shown in many instances that extracellular vesicles can contain small RNAs that exert distinct functions during infection, especially in regard to plant fungal pathogens (46, 47). We believe that a combination of factors is likely involved in the antifungal effect of extracellular vesicles.

Part of the explanation may also come from similarities between neutrophilic granules and extracellular vesicles. Granules and extracellular vesicles are distinct cellular features, but do share some biogenesis pathways and cargo molecules. For example, the azurophilic granules contain myeloperoxidase to aid in pathogen killing in the endocytic pathway via fusion of the granules to pathogen-containing phagosomes (48), but myeloperoxidase is also released in a CD63-negative population of extracellular vesicles known as microvesicles (49). Based on the proteomics analysis of this study, as well as our previous work (10), it seems possible that microvesicle-cargo proteins like myeloperoxidase contribute at least in part to the antifungal activity. However, the presence of myeloperoxidase in spontaneously released extracellular vesicles and the lack of susceptibility of patients lacking myeloperoxidase to fungal infections (50) hint that other factors are also involved. Besides, the shared marker protein CD63 nicely highlights another link between granules and extracellular vesicles, the shared biogenesis from multivesicular bodies of azurophilic granules and the extracellular vesicle subset called exosomes (51).

In conclusion our results suggest that DMF-differentiated PLB-985 cells can be used as a model to study aspects of the interaction of the human pathogenic fungus *A. fumigatus* with neutrophilic granulocytes. Although they will never substitute for all experiments with neutrophils, we do believe that this system will serve as a useful tool for the genetic dissection of *A. fumigatus* pathogenesis in the future.

## ACKNOWLEDGEMENTS

We would like to thank Johannes Wagener for providing the *A. fumigatus* mitochondrial GFP reporter strain. We also thank Natascha Wilker for excellent technical assistance. The work presented here was generously supported by the Deutsche Forschungsgemeinschaft (DFG)-funded Collaborative Research Center/Transregio FungiNet 124 ‘Pathogenic fungi and their human host: Networks of Interaction’ (210879364, project A1 and Z2), the DFG-funded ANR project (316898429, project AfuInf), and by the Federal Ministry for Education and Research (BMBF: https://www.bmbf.de/), Germany, Project FKZ 01K12012 “RFIN – RNA-Biologie von Pilzinfektionen.” The authors declare no conflicts of interest.

## SUPPLEMENTAL FIGURES

**FIG S1.**
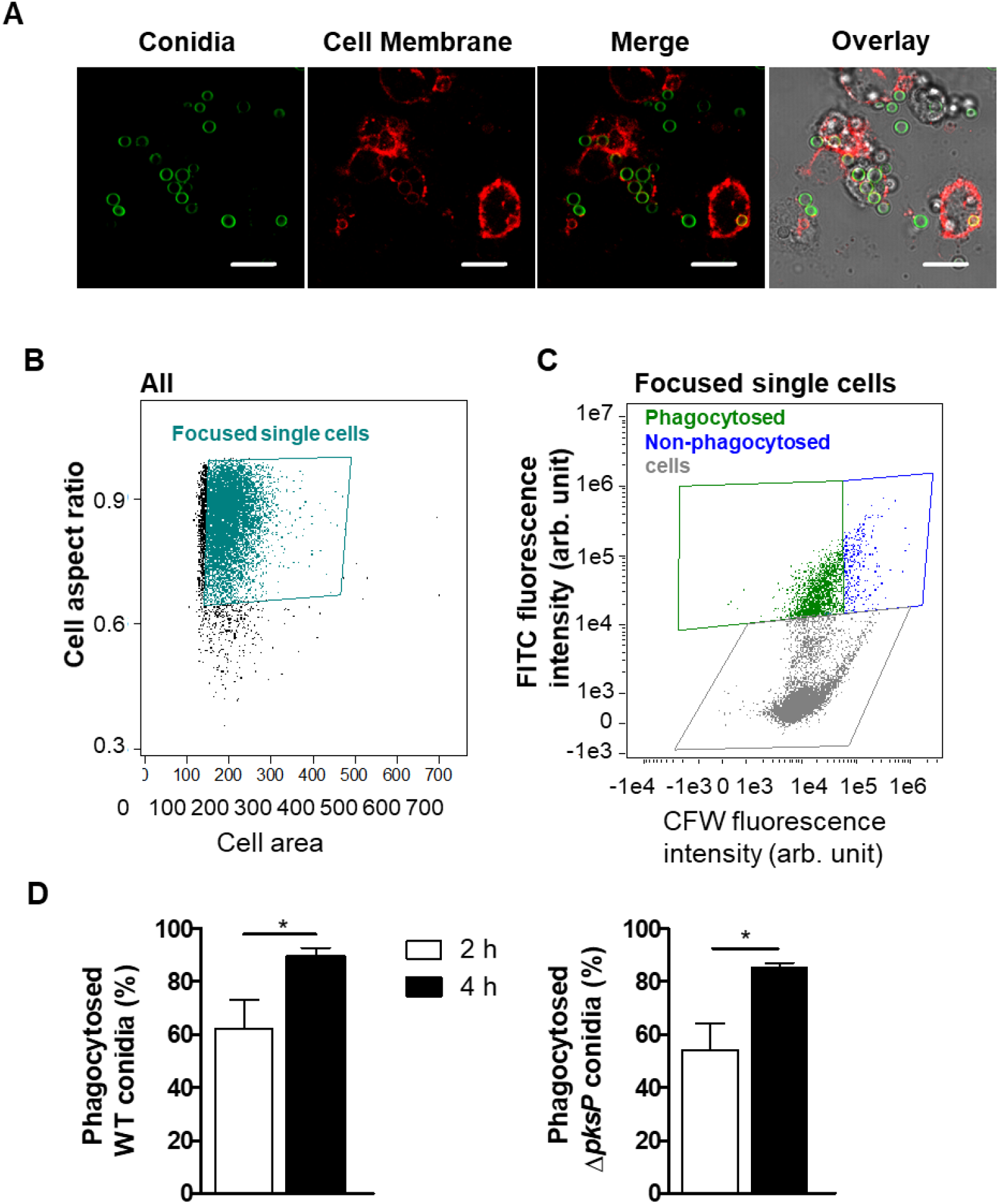
Primary human neutrophils phagocytose wild-type conidia. **(A)** Confocal microscopy images of wild-type conidia (CFW-stained; false-colored green) phagocytosed by primary human PMNs after 2 hours co-incubation. Membranes were stained with Cell Mask (red) to indicate cell membranes. Images are representative of at least two independent biological experiments. Scale bars are 10 µm. **(B and C)** indicate the gating strategy for determining phagocytosis rates of dPLB cells and PMNs using imaging flow cytometry. **(B)** Single cell populations were selected based on their area and aspect ratio in the bright field channel. **(C)** Phagocytosed conidia were gated based on their higher FITC fluorescence intensity and low CFW fluorescence intensity. Cells without conidia had low FITC fluorescence intensity. The data subject to analysis included only cells in focus and were pre-gated during data acquisition based on their high root mean square gradient in the bright field channel. **(D)** Quantification of phagocytosis by primary human PMNs using the IDEAs software on 5,000 cells per condition with either wild-type or Δ*pksP* conidia from four independent experiments.

**FIG S2.**
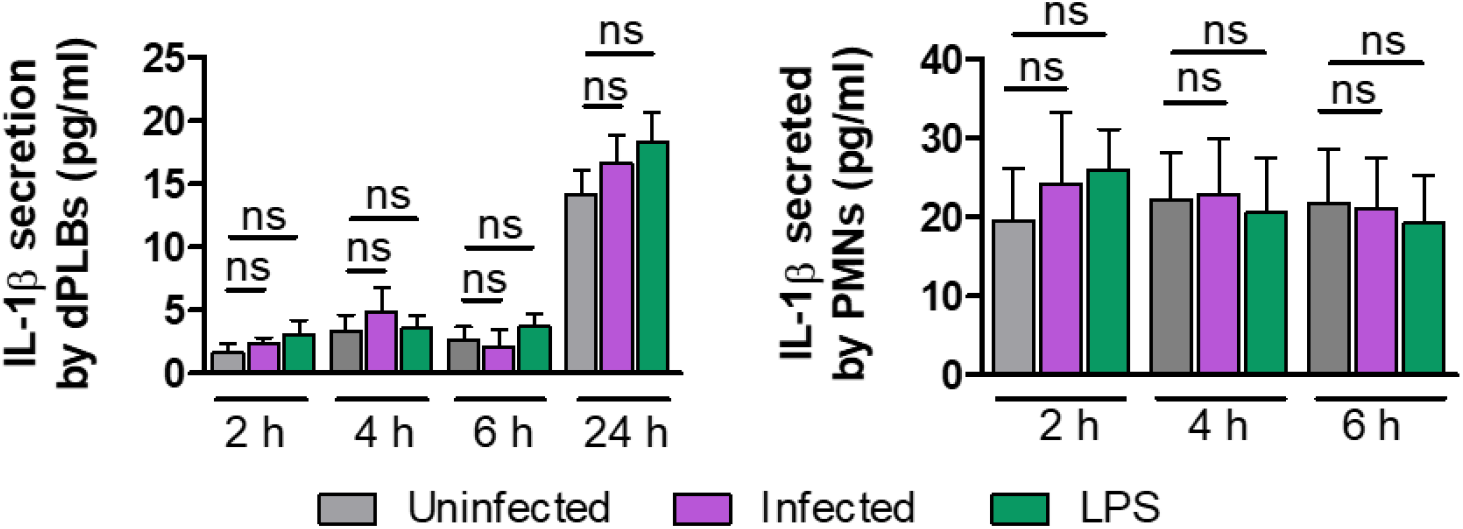
Cytokine production by dPLB cells and PMNs. ELISA detection of the IL-1β cytokine released from infected dPLBs or primary human PMNs at different time points. Lipopolysaccharide (LPS) was included as a comparison to a bacterial stimulus. Data are presented as mean ±SD, from six biological replicates; * = p < 0.05.

**FIG S3.**
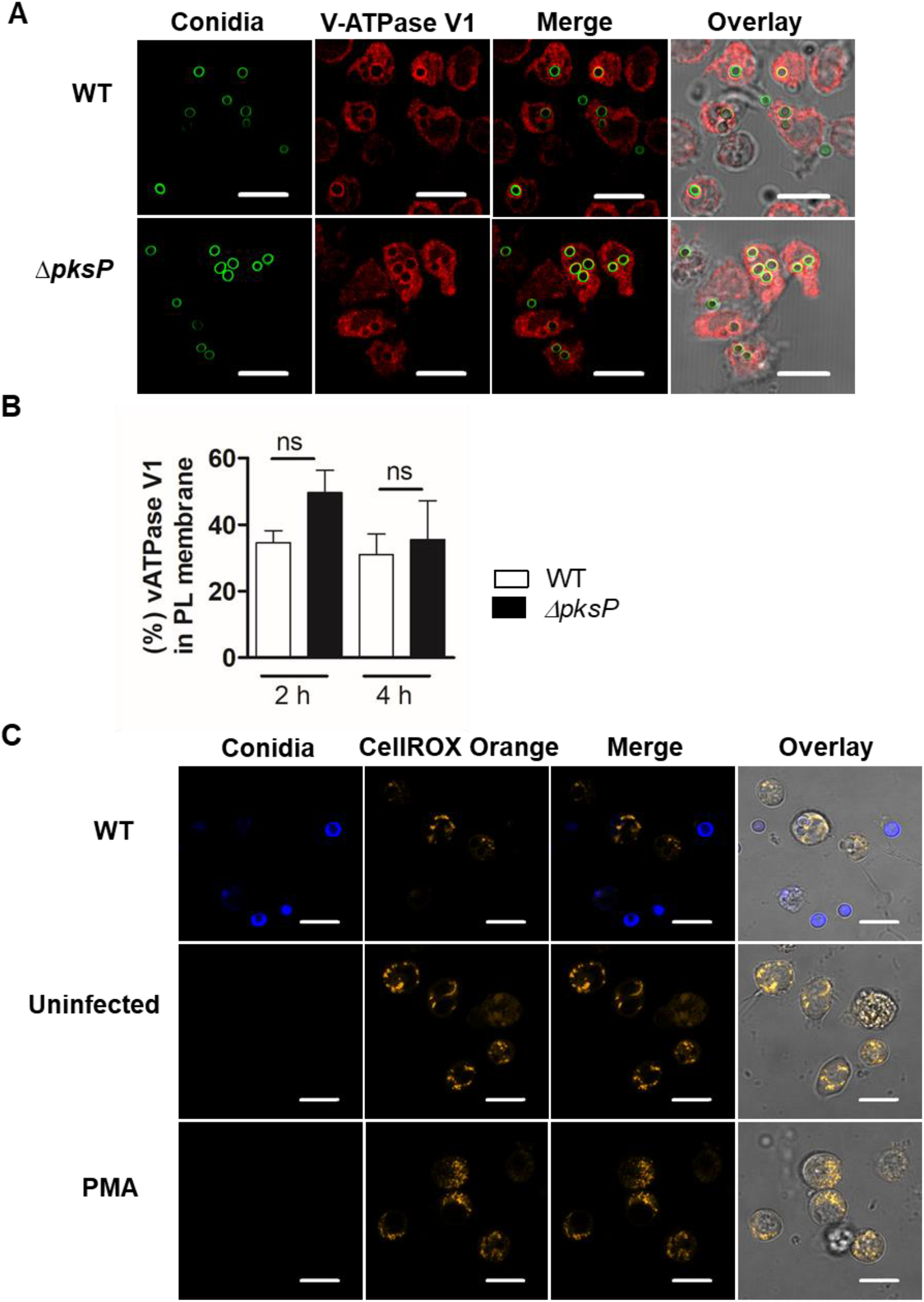
V-ATPase V1 localization to phagolysosomal membranes and ROS production by dPLB cells. **(A)** Immunofluorescence staining for V-ATPase V1 (red) in dPLB cells after infection with wild-type or Δ*pksP c*onidia (FITC labeled; green) for 4 hours. Images are representative of three biological replicates. Scale bars are 10 µm. **(B)** Quantification of V-ATPase V1 positive phagolysosomes after 4 hours of infection with wild-type or Δ*pksP* conidia. Graphs are from three biological replicates. **(C)** Confocal microscopy images of dPLB cells infected with wild-type conidia (CFW-stained; blue), left uninfected, or treated with PMA as a positive control and stained for detection of reactive oxygen species using CellROX Orange stain.

**FIG S4.**
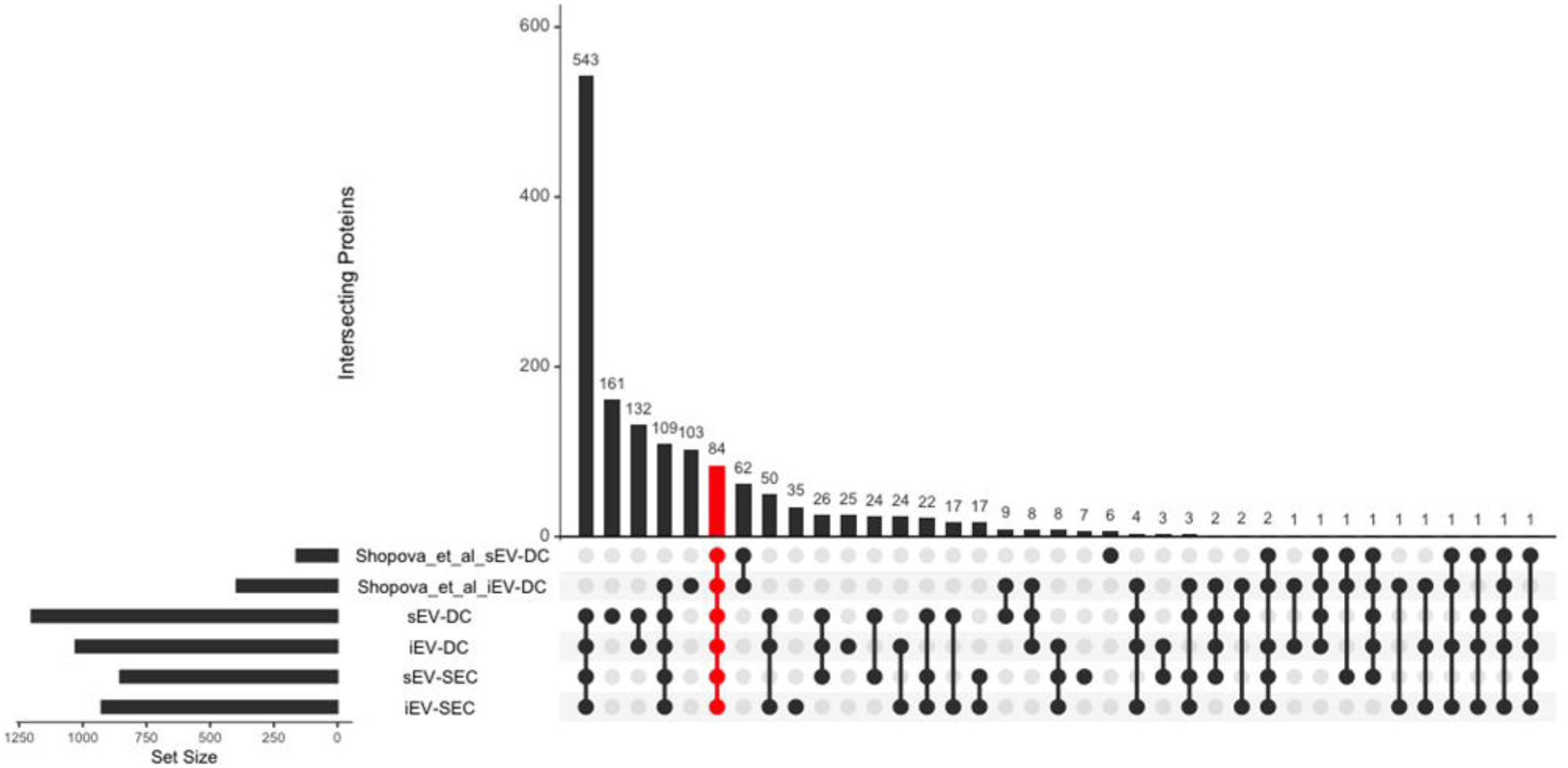
Intersection of proteomics data with previously reported neutrophil-derived EV proteomes. LC-MS/MS proteomics analysis was performed on extracellular vesicles isolated from dPLB cells using a centrifugation-based approach (DC) or a size-exclusion chromatography-based approach (SEC) in the presence or absence of infection with opsonized *A. fumigatus* conidia and compared to data from (10). Proteins that were identified in this study were compared to proteins identified by a similar approach using label-free quantification from Shopova et al. (10). Proteins had to be found in at least two replicate of a given sample to be included in the UpSetR analysis. The red bar indicates proteins that were found in all six samples.

**FIG S5.**
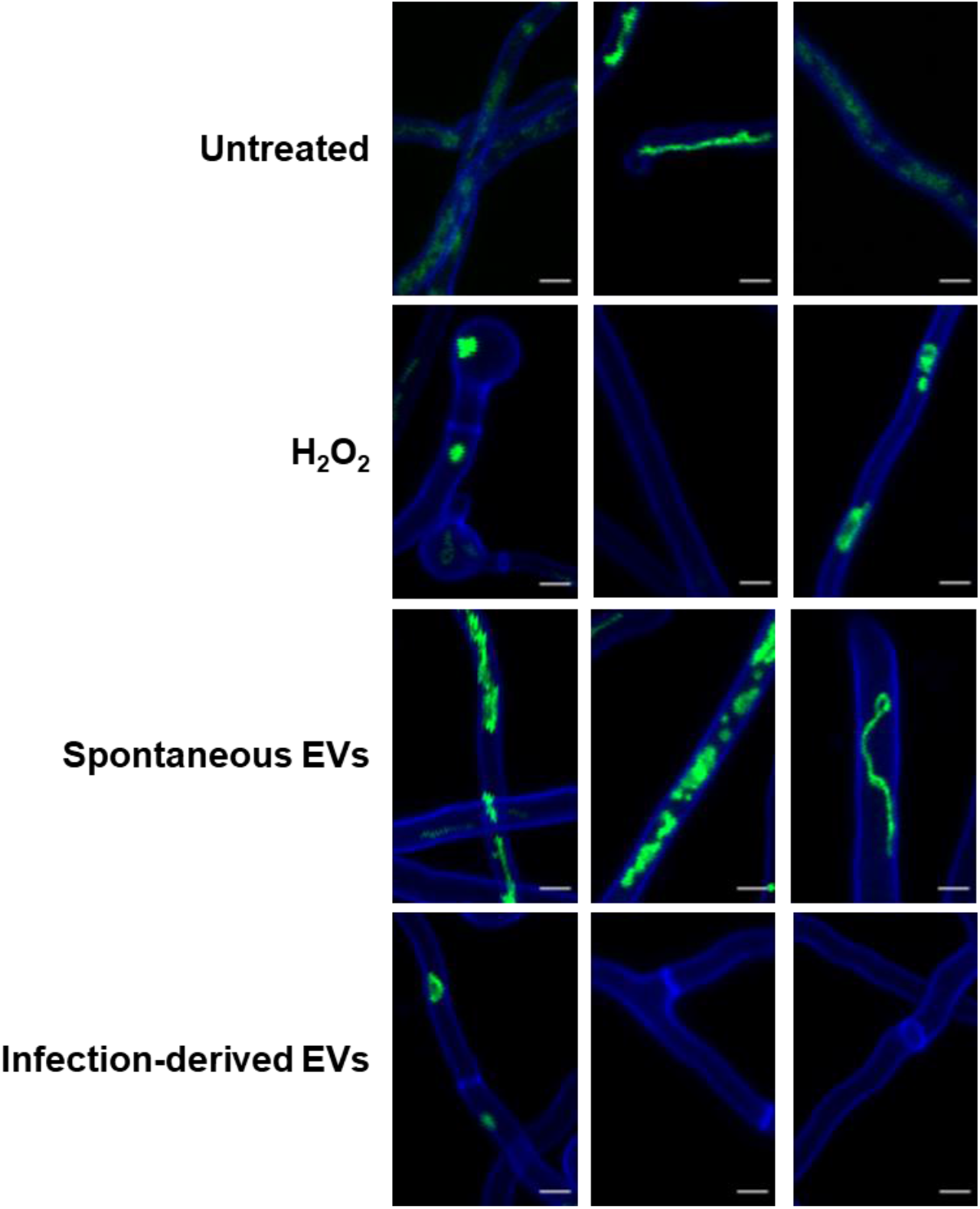
Representative images of dPLB-derived extracellular vesicles effects on *A. fumigatus* hyphae. *A. fumigatus* strain AfS35 contains plasmid pJW103, which expresses a mitochondrial GFP reporter (green). The strain was grown for 6 hours and then stained with calcofluor white (blue) and incubated overnight with spontaneously released extracellular vesicles or infection-derived extracellular vesicles from dPLBs. Additional representative images are included to show extent of variability that occurs in regard to extracellular vesicle killing experiments shown in **Fig. 6**. As a control, untreated hyphae and hyphae treated with 3 mM H_2_O_2_ to induce cell death are included. An intact mitochondrial network is shown by a filamentous network, whereas a disrupted network is shown by fragmentation or the lack of green signal. Scale bars are 5 µm.

## SUPPLEMENTAL TABLES

**Table S1.**
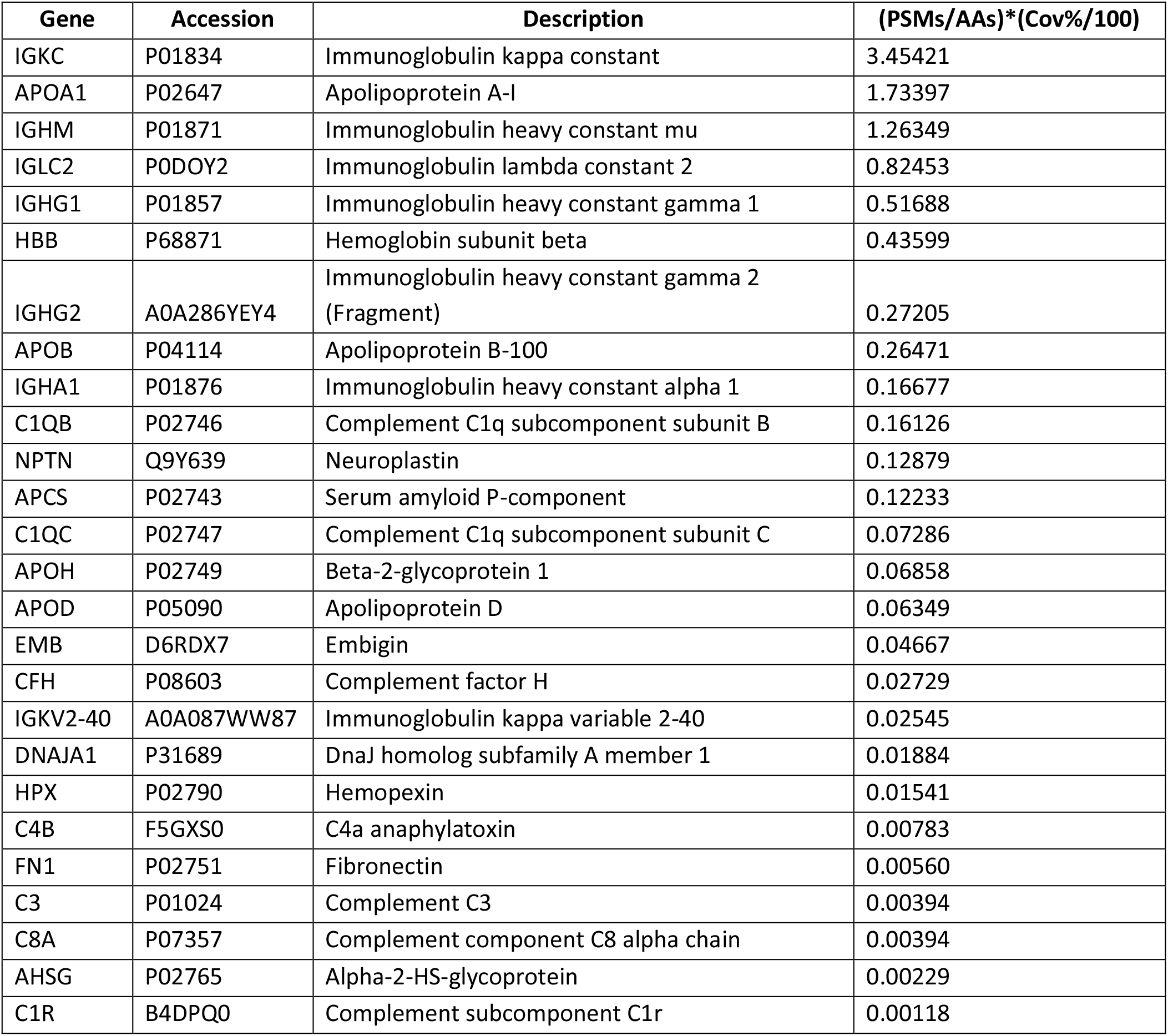
Proteins identified only in infection-derived extracellular vesicles (≥2 replicates) by both isolation methods. Abbreviations: PSMs, peptide spectrum matches; AAs, amino acids; Cov%, percent coverage of mapped peptides.

## SUPPLEMENTAL DATASETS

**Dataset S1. Proteomics data of extracellular vesicles produced in response to *A. fumigatus* infection by dPLB cells**.

